# Glucose Production and Glucose Disposal Integrated by an Intrinsic Inverse Coupling in Vivo: hypothesis of inter-organ reflex on glucose regulation

**DOI:** 10.1101/2020.03.25.009035

**Authors:** Nan Liu, Wei Zou, Ying Xing, Xi Zhang, Bin Song, Chen Wang

**Author notes:** correspondence to Chen Wang, or.

## Abstract

Glucose production (GP) and glucose disposal (Rd) are two decisive and fundamental parameters in glucose turnover and in glucose homeostasis regulation. In conventional theory, GP and Rd were responsive to regulatory factors respectively and independently of each other. Even though GP and Rd responded in reverse to insulin, GP for suppression and Rd for elevation, these inverse alterations used to be attributed to insulin multiple functions both on hepatic GP, directly or indirectly, and on whole-body glucose Rd. However, in the present study, we found GP and Rd were inversely coupled intrinsically no matter which side was the target of insulin by comparison of Rd and GP data pairs between peripheral vein insulin infusion protocol and portal vein insulin infusion protocol in rats. Furthermore, neither circulating NEFA nor HFD induced resistance broke the GP-Rd inverse coupling, but both of them reduced the responses of both GP and Rd to insulin. In conclusion, we provide the evidence that GP and Rd are two coupled parameters in vivo and they alter in reverse simultaneously, the mechanism under which needs further investigation but we tend to believe an inter-organ neural reflex was involved.

---

The blood glucose homeostasis is regulated very well no matter in physical exercise or during sleep and no matter in fed or in fasting. This homeostasis regulation needs the coordinated works of different organs, in which glucose production (GP) of liver and glucose disposal (Rd) of whole-body tissue are the most important two sides of the glycemia balance. Various hormones, insulin, epinephrine, and glucagon etc. are involved in the integrated coordination of regulating the blood glucose balance. Insulin orchestrates both the whole-body glucose disposal and glucose production to slant the balance towards blood glucose decreasing.

Following acute administration of insulin, the GP would be inhibited and the Rd would also be enhanced, which two effects both contribute to lower blood glucose levels. Some solid evidences indicated that insulin directly inhibited GP^1^, while other unequivocal studies showed that the effect of insulin on GP was indirect^2^. The direct effect model acted as these two effects, GP suppression and Rd elevation, came respectively from insulin’s two arms independently, one for liver to suppress of GP and the other one for peripheral tissues to elevate Rd. The indirect effect model otherwise indicated that the inhibition of GP was partially or totally secondary to insulin’s effects on peripheral tissues (i.e. the inhibition of adipose lipolysis)^2^ and the insulin signal in liver was even dispensable for suppression of GP^3^. Evidences showed that severe impairment in liver insulin signaling failed to alter hepatic glucose production in conscious mice^4^. These results looked as fortunate as one stone two birds. Only one insulin stone acts on peripheral tissue will get two birds as Rd promotion and GP inhibition. On the other hand, if we threw one insulin stone to liver, could we also get the same two birds as GP inhibition and Rd promotion? It was reported that a muscle-specific insulin receptor knockout model could exhibit normal glucose tolerance^5^. Moreover, if both adipose and muscle insulin action were impaired, the blood glucose level was still normal although with insulin resistance^6^. More intriguingly, restoration of liver specific insulin signaling in the whole-body insulin receptor knockout mice could restore the whole-body Rd^7^. Therefore, we assumed that the action of insulin on hepatic GP could also be transmitted to wholebody Rd, which constituted a bidirectional coupled reactions of GP and Rd in glucose regulation independent of insulin. These bidirectional coupled reactions gave us a clue that there might be an integrated relationship between GP and Rd throughout the whole body, independent of insulin. We hypothesized that there was an intrinsic inverse coupling between GP and Rd in vivo, i.e. if GP altered for any reason, the whole-body Rd would alter in reverse at the same time, and vice versa. The most common form of the coupling is GP falling coupled with Rd rise, which used to be masked under the shadow of insulin’s multifunction on both hepatic glucose production and peripheral glucose disposal.

To examine whether GP and Rd were integrated by an intrinsic inverse coupling independent of insulin, we used two different hyperinsulinemic-euglycemic clamps in rats (Peri and Port). Under pancreatic clamp with somatostatin (3μg/kg·min) and basal glucagon infusion (1.5ng/kg·min), we used different insulin infusion rates to get pairs of Rd-GP data. In peripheral protocol (Peri), insulin was infused through peripheral vein of rats and glucose infusion rate varied to keep euglycemia, in which the direct insulin actions preferred peripheral tissues than the liver^8^ as the peripheral artery plasma insulin concentration is higher than in portal vein (Fig.1A). And the indirect effect of insulin on GP suppression was supposed to be mediated by lower level of circulating NEFA from reduced adipose lipolysis^9,10^. After normalized by basal level glucose turnover rate, the Rd-GP data pairs at 90 min during clamp showed an inverse coupling relationship (Fig.1B, C, R^2^=0.97). To exclude the explanation that the GP-Rd coupling was only occasional and attributed to the direct and indirect multifunction of insulin, we also did the portal protocol (Port), in which insulin was infused through portal vein system of rats as a model of insulin direct effect preferred liver^11,12^. In this model, the portal vein insulin concentration was much higher than in peripheral artery (Fig.1A) and it was reported that the direct action of insulin on GP suppression was dominant in this protocol^13^. In this situation, we also got Rd-GP data pairs according to different insulin infusion rate (Fig.1B, and Fig.1C, R^2^=0.95). Intriguingly, the response of Rd-GP data pair to insulin infusion was similar between the two protocols, the only difference was higher insulin infusion rates in Port protocol. There was no predilection for GP suppression in Port group or for Rd elevation in Peri group (Fig.1B,C) despite of the significant polarized of insulin distribution between peripheral artery and portal vein of the two groups (Fig.1A). On the contrary, Rd elevation was significant higher in Port group coupled with the lower of GP than in Peri group at the same artery insulin concentrations (Fig.1D), and GP suppression was deeper in Peri group coupled with the higher Rd than in Port group at the same portal vein insulin concentrations (Fig.1E). The results also showed that GP suppression was more responsive to insulin concentration increasing at a lower range of insulin concentrations but Rd elevation was more responsive to the increasing at a higher range of insulin concentrations in both groups. Furthermore, each replicates of the Rd and GP alterations in two groups fit very close lines of regression (Fig.1C, R_Peri_^2^=0.97, R_Port_^2^=0.95) and fit very well the combination curve of regression (Fig.1F, R^2^=0.96). In conclusion, we got indistinguishable coupled Rd-GP pairs between the Peri and the Port protocols (Fig.1B,C,D,E,F) despite the polarized imbalance of insulin distribution in the two groups (Fig.1A). During the experiment, plasma lactate, glucagon, epinephrine, norepinephrine and corticosterone levels were not different between the two groups. The only explanation was that GP and Rd altered as a whole together undivided but not as occasionally by multifunction of insulin on both GP and Rd. Based on these results, we confirmed the existence of inverse coupling of GP-Rd. Therefore, the action of insulin would not only pass from extrahepatic tissues to liver to suppress GP but also would pass from liver to extrahepatic tissues to elevate Rd because of this GP-Rd coupling.

In our opinion, the correlation of Rd and GP seems like a seesaw coupling. For example, if we elevated a seesaw on the right side (as Rd) to Position 1, the left side (as GP) would get to an suppressed Position 2. Next time, if we went to left side of the same seesaw and suppressed the left side (as GP) to the same Position 2, the right side (as Rd) would with no doubt increase to the same Position 1 (Fig.1G). You couldn’t distinguish what side we were manipulating the seesaw if observing merely the slope angle of the seesaw. The results also imply that any acute suppressive factor for GP will synchronously enhance whole-body glucose disposal, and any intervention to elevate whole-body glucose disposal will necessarily reduce glucose production accordingly. The whole-body glucose disposal Rd in the present study included the glucose uptake by muscle, adipose tissue, brain and liver etc., and the Rd elevation was tightly integrated with GP suppression but the glucose uptake contribution to Rd between different tissues could be heterogeneous and dynamic under different conditions, i.e. if insulin infused through portal vein system, the glucose uptake was equally divided into liver and muscles, but if insulin infused into peripheral vein system, the glucose uptake by muscles was four-fold greater than by liver^14^. In this study, we only testified the part of GP suppression with Rd elevation in the GP-Rd inverse coupling as the other part of GP elevation with Rd suppression, i.e. administration of glucagon, would inevitably increase the blood glucose level and steady state could not reached until GP equal to Rd again. And the glucose level elevation would shift the GP-Rd coupling regression curve to right (Fig.1H). With the right shift of the curve, we could probably see the GP elevation simultaneously accompanied by Rd elevation, which was like that the both two sides of seesaw elevated at the same time if the axis of the seesaw was elevated, but the seesaw coupling per se was still intact (Fig.1I). Similarly, during long time fasting, the plasma glucose level would fall slowly which would shift the GP-Rd coupling curve to left (Fig.1J), and in this condition, Rd reduction with GP suppression could appear at the same time but that didn’t mean the coupling broken.

**Figure 1.**
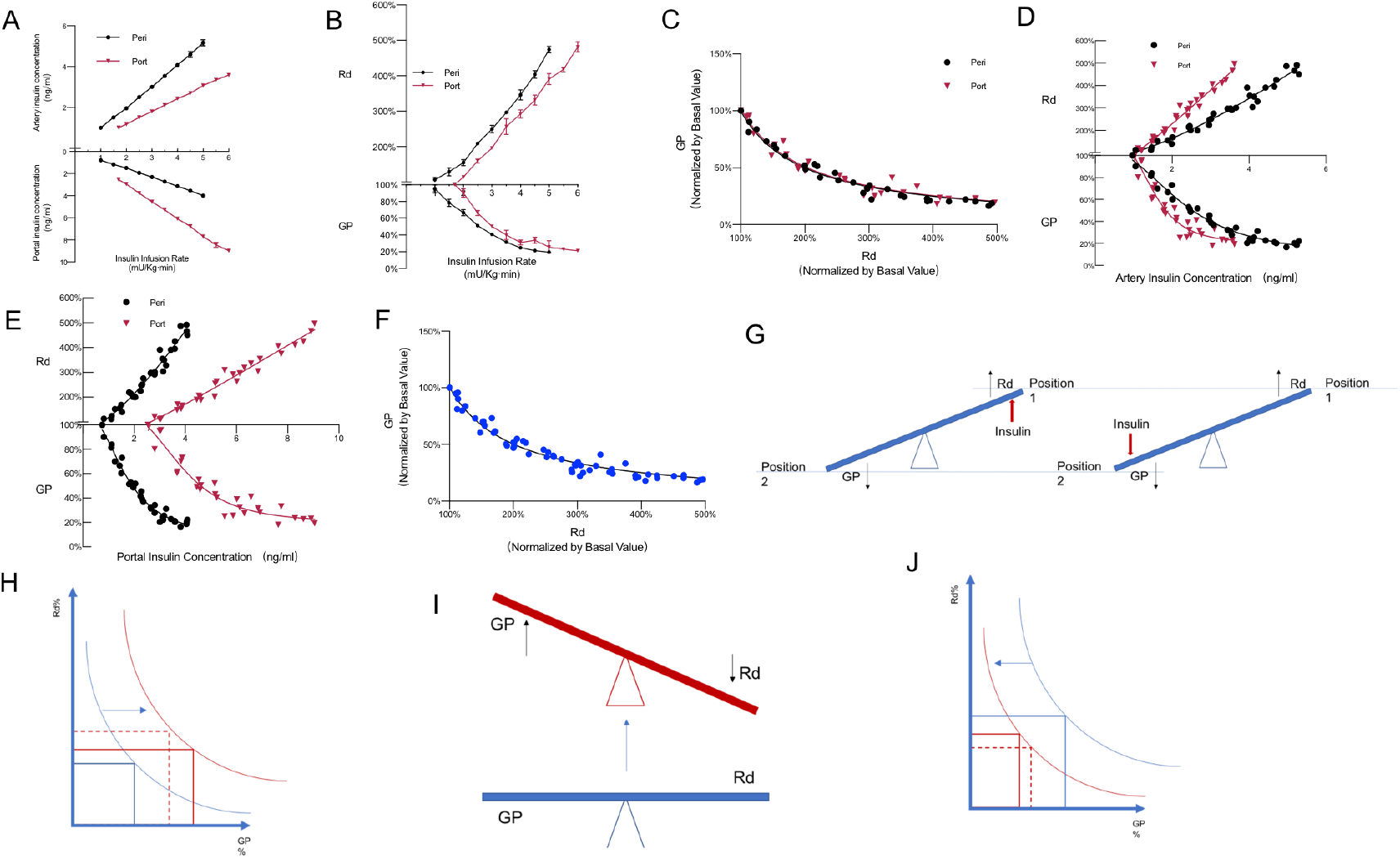
The correlation of Rd and GP was coupled independently of insulin distribution drift. A. Insulin concentration and distribution imbalance in Peripheral protocol(Peri) and portal protocol(Port).Different animals were used in different infusion rates. Data are mean ± SEM. Difference were significant(P<0.01) from 2 mU/kg·min insulin infusion on both artery concentration and portal vein concentration. B. Insulin infusion on glucose disposal(Rd) and glucose production (GP). C. GP and Rd correlation. D. Correlation between artery insulin concentration and Rd, GP. E. Correlation between portal vein insulin concentration and Rd, GP. F. Correlation of Rd and GP combined from both groups. G. GP and Rd seem like the two sides of a seesaw coupled as a whole. H. Glucose level elevation would shift the curve to right with both GP and Rd increasing. I. The seesaw elevation with both sides increasing. J. Glucose level falling would shift the curve to left with both GP and Rd decreasing.

As excess circulating NEFA, derived from excess peripheral lipolysis, drove the higher rates of GP^15^, and NEFA levels were lower in Peri group than in Port group at a same insulin infusion rates in the present study (Fig.2A), it was plausible to attribute the hepatic GP suppression to the lower circulating NEFA alternatively rather than the intrinsic GP-Rd coupling. To test if NEFA was the bridge of the GP-Rd coupling, we repeated the Peri protocol with 20% intralipid infusion with heparin to maintain plasma NEFA to two higher levels (0.4ml/kg·h, Low lipid and 5ml/kg·h, Hi lipid, Fig.2B). Results showed that the higher levels of NEFA would need the higher insulin infusion rates to get comparable Rd-GP data pairs and the GP-Rd coupling were the lower responsive to insulin. However, the suppression of GP and the elevation of Rd by insulin were both blunted by higher circulating NEFA (Fig.2C,D), and the coupled GP-Rd correlation did not change (Fig.2E,F). The combination of saline group with Hi-Lipid group and Low-Lipid group fits very well the GP-Rd regression curve (Fig.2G, R^2^=0.95). These results indicated that the GP-Rd coupling was not dependent on circulating NEFA level but the higher of NEFA level could act effects on the GP-Rd coupling as a whole to resist the action of insulin on both sides (Fig.2H).

**Figure 2.**
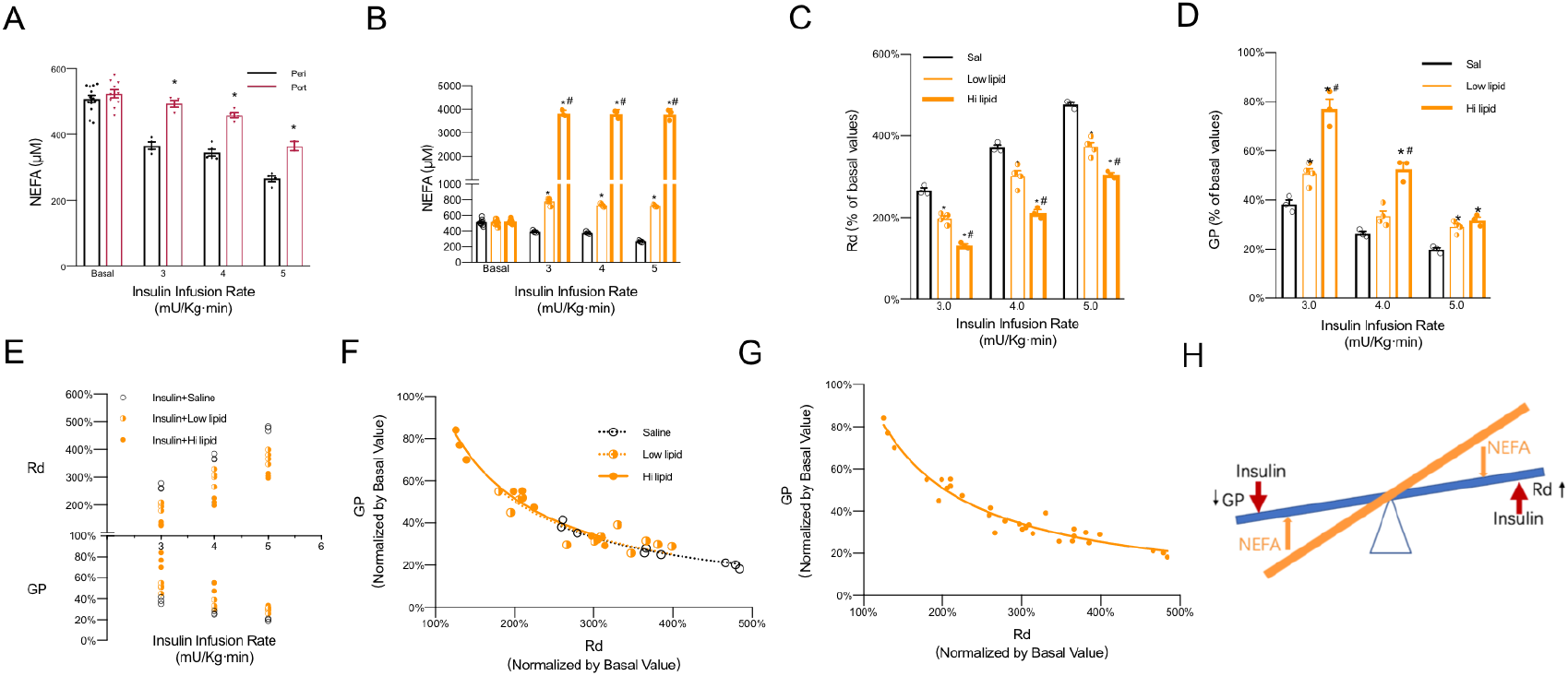
The coupled Rd and GP were independent of circulating NEFA. A. Circulating NEFA difference between Peri and Port protocols. * P<0.01 compared to Peri. B. Low and high lipid infusion elevated NEFA to different levels during the clamp. * P<0.01 compared to saline group, # P<0.01 compared to low lipid group. Different rats were used in different insulin infusion clamps. C.NEFA dampened the Rd response to insulin. * P<0.01 compared to saline group, # P<0.01 compared to low lipid group. D. NEFA dampened the GP response to insulin. * P<0.01 compared to saline group, # P<0.01 compared to low lipid group. E. Each replicates of Rd-GP responses to insulin in three groups. F. GP and Rd correlation regression curves in three groups. G. GP and Rd correlation regression curve combined. H. NEFA effect on GP-Rd coupling to resist insulin action.

In order to examine whether this coupling was different in high-fat-diet (HFD) induced insulin resistant model, we repeated the Peri and Port hyperinsulinemic-euglycemic protocol in one week HFD fed rats (HFD-Peri, HFD-Port) and in normal diet rats (ND-Peri). The basal glucose turnover rate was significantly higher in HFD groups with a higher basal plasma glucose level than in ND groups (Fig.3A, B). During clamps, the insulin infusion rates in HFD groups were higher than in ND groups when Rd-GP data pairs got to a close levels of the ND group, and HFD-Port needed higher insulin infusion rates than HFD-Peri (Fig.3C), which was consistent with Peri and Port groups in normal rats in Fig.1A. Because of our clamping plasma glucose level from ~9mM to 6mM in HFD groups, we found that this significant decreasing of plasma glucose concentration induced a significant reduction in both Rd and GP in HFD groups to about 66% of basal value with no need of glucose infusion to maintain the 6mM glucose level at lower insulin infusion rates (~3 mU/kg·min in HFD-Peri, ~5 mU/kg·min in HFD-Port) (Fig.3C), which was consistence with our hypothesis of curves shift to left in the situation of glucose falling (Fig.1J,3D). However, the HFD induced insulin resistance didn’t break the GP-Rd inverse coupling despite GP and Rd were both synchronously suppressed by insulin infusion as the regression curve fit well if we combined the HFD-Peri and HFD-Port into HFD-comb (Fig.3D, R^2^=0.88). The deviation of the two curves between ND-Peri and HFD-Comb was attributable to the glucose level decreasing from ~9mM to our clamping 6mM. Based on these results, it was concluded that the GP-Rd inverse coupling worked well in HFD induced rat insulin resistant model. These results also implied that there was no hepatic-specific or extrahepatic-tissue-specific insulin resistance in vivo in HFD rat model because liver and other tissues were coupled together as a whole in response to insulin, which was consistence with the reports that impaired Rd in adipose tissue or in muscle would impair the hepatic GP response to insulin independent of circulating NEFA, triglycerides or leptin^16,17^.

**Figure 3.**
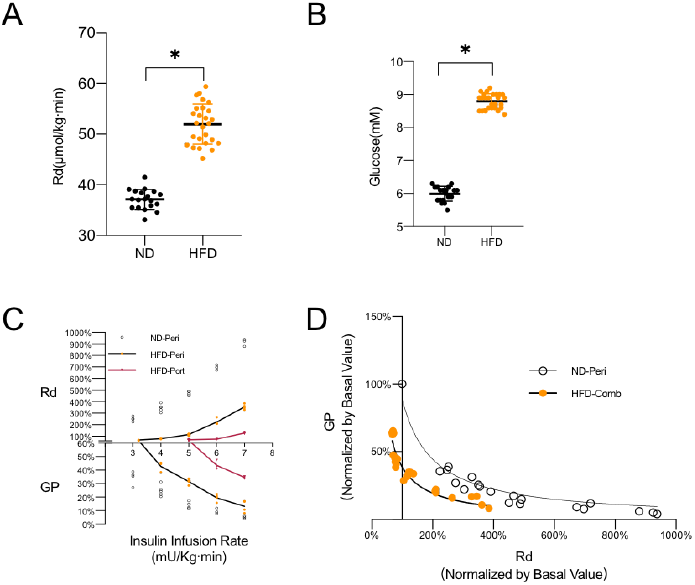
HFD effects on GP-Rd coupling in rats. A. Basal glucose turnover rates in HFD rats were higher than normal rats. * P<0.01. B. Basal glycemia in HFD rats were higher than normal rats. * P<0.01. C. Clamp glycemia to 6mM with insulin infusion induced a significant decreasing of glucose turnover to about 66% with no need of additional glucose infusion in the both HFD groups. D. the deviation of the two GP-Rd correlation regression curves in normal rats and in HFD rats under same level of euglycemic clamps.

In the present study, we didn’t distinguish the contribution from gluconeogenesis flux or the glycogenolysis flux to GP but we believed the contribution proportion would vary according to different situations while the gross total GP was coupled with whole-body Rd. The reason why the portal protocol needed higher insulin infusion rates could be accounted for by the hepatic first-pass degradation of insulin but we couldn’t exclude the possibility of that the resistance was different if insulin acted on different side of the GP-Rd coupling. The mechanism of the GP-Rd inverse coupling is not clear but we tended to believe that it is mediated by the reflex of neural system because the coupling is real-time responded and synchronized simultaneously, in which the hypothalamus might be involved^15,18,19,20^ or not be necessarily involved^21^. In conventional theory, the correlation of hepatic GP and whole-body Rd was connected and orchestrated by hormones i.e. insulin, glucagon, epinephrine, corticosterone etc. or by metabolites i.e. NEFA, glycerol, lactate etc., which were all humoral regulation factors. In the present study, we found that there might be an additional neural regulation mechanism between the inter-organ GP-Rd coupling. We speculated the reflex arc was intact in lower spinal cord just like the tendon reflex arc if hypothalamus was not involved. ATP-sensitive potassium (K_ATP_) channels was reported as a pivotal element connecting the neural system to both glucose disposal and glucose production^22,23^, so we speculated that the glucose disposal stimulated K_ATP_ channels on neural fiber ends embedded in peripheral tissues and then the signals transmitted to liver and suppressed GP. On the other hand, the suppression of GP would also stimulate K_ATP_ channels on neural fiber ends embedded in liver then transmitted the signals to whole-body tissues to enhance Rd. In conclusion, the whole-body Rd was integrated with the endogenous GP as GP-Rd inverse coupling to orchestrate a coordinated responses for blood glucose homeostasis regulation, the mechanism under which needs further investigations to confirm.

## Methods

We studied 14-week-old male Sprague Dawley rats fed with normal chow or high fat diet for one week. Animal preparation was performed as previously described^24^. Insulin was infused through the jugular vein catheter in the peripheral protocol, and catheters were placed in the ileal vein for insulin infusion in the portal protocol. Blood samples were collected from left common carotid artery catheter or portal vein as needed. To calculate GP and Rd, [3-^3^H] glucose was given at 20 μ Ci in a bolus 120min before clamp and followed by continuous infusion at 0.2 μ Ci/min and calculated by Steele’s equation^25^. Different rats were used in different rates of insulin infusion clamps. Data are expressed as mean ± SEM. Statistical analyses were performed using GraphPad Prism 8. *P* values were calculated by one-way or two-way ANOVA or by unpaired Student’s *t*-test where appropriate.

## ACKNOWLEDGMEENTS

These studies were supported by grants from National Nature Science Foundation of China (30900611 to C.W., 81101959 to W.Z., and 30901336 to Y.X.) and supported by Young Talent Fund of University Association for Science and Technology in Shaanxi, China (20170404 to N.L.)

## AUTHOR CONTRIBUTIONS

N.L. conducted and designed experiments, performed data analyses and wrote the manuscript. W.Z., X.Z. and B.S. assisted with experiments in animal preparation and surgery. Y.X. and C.W. designed experiments and edited the manuscript. C.W. proposed the hypothesis and adapted the protocols and supervised the study.

## References

1. Edgerton, D. S. et al. Insulin’ s direct hepatic effect explains the inhibition of glucose production caused by insulin secretion. JCI Insight 2, e91863 (2017).

2. Perry, R. J. et al. Hepatic acetyl CoA links adipose tissue inflammation to hepatic insulin resistance and type 2 diabetes. cell 160, 745–758 (2015).

3. Titchenell, P. M. et al. Direct Hepatocyte Insulin Signaling Is Required for Lipogenesis but Is Dispensable for the Suppression of Glucose Production. Cell Metab. 23, 1154–1166 (2016).

4. Buettner, C. et al. Severe impairment in liver insulin signaling fails to alter hepatic insulin action in conscious mice. J. CHn. Invest. 115, 1306–1313 (2005).

5. Bruning, J. C. et al. A muscle-specific insulin receptor knockout exhibits features of the metabolic syndrome of NIDDM without altering glucose tolerance. Mol. Cell 2, 559–569 (1998).

6. Lauro, D. et al. Impaired glucose tolerance in mice with a targeted impairment of insulin action in muscle and adipose tissue. Nat. Genet. 20, 294–298 (1998).

7. Okamoto, H., Obici, S., Accili, D. & Rossetti, L. Restoration of liver insulin signaling in *Insr* knockout mice fails to normalize hepatic insulin action. J. Clin. Invest. 115, 1314–1322 (2005).

8. Farmer, T. D. et al. Comparison of the physiological relevance of systemic vs. portal insulin delivery to evaluate whole body glucose flux during an insulin clamp. Am. J. Physiol.-Endocrinol. Metab. 308, E206–E222 (2015).

9. Sindelar, D. K. et al. The role of fatty acids in mediating the effects of peripheral insulin on hepatic glucose production in the conscious dog. Diabetes 46, 187–196 (1997).

10. Lewis, G. F., Zinman, B., Groenewoud, Y., Vranic, M. & Giacca, A. Hepatic glucose production is regulated both by direct hepatic and extrahepatic effects of insulin in humans. Diabetes 45, 454–462 (1996).

11. Sindelar, D. K., Balcom, J. H., Chu, C. A., Neal, D. W. & Cherrington, A. D. A comparison of the effects of selective increases in peripheral or portal insulin on hepatic glucose production in the conscious dog. Diabetes 45, 1594–1604 (1996).

12. An, J. et al. Hepatic expression of malonyl-CoA decarboxylase reverses muscle, liver and whole-animal insulin resistance. Nat. Med. 10, 268–274 (2004).

13. Edgerton, D. S. et al. Insulin’ s direct effects on the liver dominate the control of hepatic glucose production. J. Clin. Invest. 116, 521–527 (2006).

14. Edgerton, D. S. et al. Targeting insulin to the liver corrects defects in glucose metabolism caused by peripheral insulin delivery. JCI Insight 5, (2019).

15. Perry, R. J. et al. Leptin reverses diabetes by suppression of the hypothalamic-pituitary-adrenal axis. Nat. Med. 20, 759–763 (2014).

16. Abel, E. D. et al. Adipose-selective targeting of the GLUT4 gene impairs insulin action in muscle and liver. Nature 409, 729–733 (2001).

17. Zisman, A. et al. Targeted disruption of the glucose transporter 4 selectively in muscle causes insulin resistance and glucose intolerance. Nat. Med. 6, 924–928 (2000).

18. Stanley, S. et al. Identification of neuronal subpopulations that project from hypothalamus to both liver and adipose tissue polysynaptically. Proc. Natl. Acad. Set. U. S. A. 107. 7024–7029 (2010).

19. Obici, S., Zhang, B. B., Karkanias, G. & Rossetti, L. Hypothalamic insulin signaling is required for inhibition of glucose production. Nat. Med. 8, 1376–1382 (2002).

20. Bentsen, M. A., Mirzadeh, Z. & Schwartz, M. W. Revisiting How the Brain Senses Glucose-And Why. Cell Metab. 29, 11–17 (2019).

21. Ramnanan, C. J., Edgerton, D. S. & Cherrington, A. D. Evidence against a physiologic role for acute changes in CNS insulin action in the rapid regulation of hepatic glucose production. Cell Metab. 15, 656–664 (2012).

22. Pocai, A. et al. Hypothalamic K(ATP) channels control hepatic glucose production. Nature 434, 1026–1031 (2005).

23. Könner, A. C. et al. Insulin action in AgRP-expressing neurons is required for suppression of hepatic glucose production. Cell Metab. 5, 438–449 (2007).

24. Shiota, M. Measurement of glucose homeostasis in vivo: combination of tracers and clamp techniques. Methods Mol Biol Clifton NJ 933, 229–253 (2012).

25. Steele, R., Wall, J. S., De Bodo, R. C. & Altszuler, N. Measurement of size and turnover rate of body glucose pool by the isotope dilution method. Am. J. Physiol. 187, 15–24 (1956).

